# A chromosome-level genome assembly of an endangered large carnivore, the wolverine, *Gulo gulo gulo* (Linnaeus, 1758)

**DOI:** 10.64898/2026.11.02.763642

**Authors:** Ole K. Tørresen, Øystein Flagstad, Oddmund Kleven, Marius A. Strand, Morten Skage, Giada Ferrari, Ave Tooming-Klunderud, Kjetill S. Jakobsen

## Abstract

We describe a chromosome-level, haplotype-resolved genome assembly from a male wolverine (*Gulo gulo gulo*). The assembly comprises two pseudo-haplotypes of 2,523.2 Mb and 2,288.4 Mb, with 95.8% and 97.8% scaffolded into 20 autosomal chromosomes and sex chromosomes (placed in haplotype one). Assembly completeness is high (BUSCO 98.2% and 88.9%), with 20,337 and 17,692 annotated protein-coding genes for haplotypes one and two, respectively. Compared to existing wolverine genomes, the current genome is more complete and contains both sex chromosomes. This reference genome will be an important resource for population studies and conservation biology of this large carnivore.

## Introduction

The wolverine (*Gulo gulo*) is a wide-ranging, cold-adapted carnivore of the Northern Hemisphere and the largest terrestrial member of the weasel family (Mustelidae) (Aubry et al., 2007; Copeland et al., 2010; Persson et al., 2023). Taxonomically, *Gulo gulo* is divided into two subspecies, *G. g. gulo* in Eurasia and *G. g. luscus* in North America (Copeland & Whitman, 2003). Occupying boreal, alpine, and tundra ecosystems across its circumpolar range, the species plays a distinctive ecological role as both predator and scavenger, linking trophic levels through carrion use and opportunistic hunting (Andrén et al., 2011; Mattisson et al., 2011; van Dijk et al., 2008). Wolverines are characterized by large home ranges, low population densities, and specialized life-history traits tied to cold environments (Aubry et al., 2007; Persson et al., 2023). These ecological features, combined with sensitivity to habitat fragmentation and climate-driven snow loss, have made the species a focal taxon in conservation biology and an indicator of environmental change in northern ecosystems (Aubry et al., 2023; Copeland et al., 2010; McKelvey et al., 2011; Ruggiero et al., 2007). Globally, the wolverine is listed as least concern in the IUCN Red List of threatened species (Abramov, 2015).

Biologically, the wolverine is notable for a suite of morphological and physiological adaptations to extreme environments, including a robust build, powerful dentition capable of processing frozen carrion, and locomotor traits suited to deep snow (Copeland et al., 2010; Thiel et al., 2019). Its behavioral ecology, marked by solitary habits, extensive dispersal, and opportunistic foraging, further distinguishes it among carnivores (Aronsson & Persson, 2018). From an evolutionary perspective, *the wolverine* occupies a key phylogenetic position within Mustelidae, with mitochondrial data placing it close to the genus *Martes* (B. Li et al., 2014). Its evolutionary history is also shaped by Pleistocene climatic fluctuations, with genetic evidence indicating persistence in northern refugia and subsequent range shifts (Krejsa et al., 2021; Zigouris et al., 2013), making it a valuable system for studying biogeography and adaptation to glacial–interglacial cycles.

In Fennoscandia, the wolverine is classified as endangered in Norway and Finland (Hyvärinen et al., 2019; Norwegian Biodiversity Information Centre, 2021) and vulnerable in Sweden (Swedish Species Information Centre, Swedish University of Agricultural Sciences, 2025). This requires a management regime balancing conservation objectives with socio-economic considerations, particularly those related to semi-domesticated reindeer husbandry and free-ranging sheep (Fisher et al., 2022; Landa et al., 1999). Monitoring programs employ a combination of non-invasive genetic sampling, snow tracking, and camera surveillance to estimate population size, reproductive success, and spatial distribution. These empirical data inform management interventions, including regulated culling and protective measures. Transboundary cooperation between the three Nordic countries is critical, given the species’ extensive range and mobility.

Genomic data can inform conservation strategies by enabling the assessment of genetic diversity, adaptive potential, and population connectivity at unprecedented resolution (Hohenlohe et al., 2021; Shafer et al., 2015). It also provides the foundation for monitoring genetic health, identifying locally adapted populations, and predicting responses to environmental change. Until recently, genomic resources for the wolverine and closely related mustelids have been sparse or fragmented. A short-read draft genome has been generated for the Eurasian wolverine (*G. g. gulo*) (Ekblom et al., 2018) and a chromosomal-level genome assembly has been generated for the North American wolverine (*G. g. luscus*) (Lok et al., 2022).

Here, we present a state-of-the-art haplotype-resolved genome assembly of the Eurasian wolverine (*G. g. gulo*) sampled from the endangered Norwegian population, generated using PacBio HiFi long-read and Hi-C sequencing data (Figure 1). This genome shows higher completeness and includes both sex chromosomes thus providing a new and improved resource for both research and conservation of this icon species of the north.

**Figure 1.**
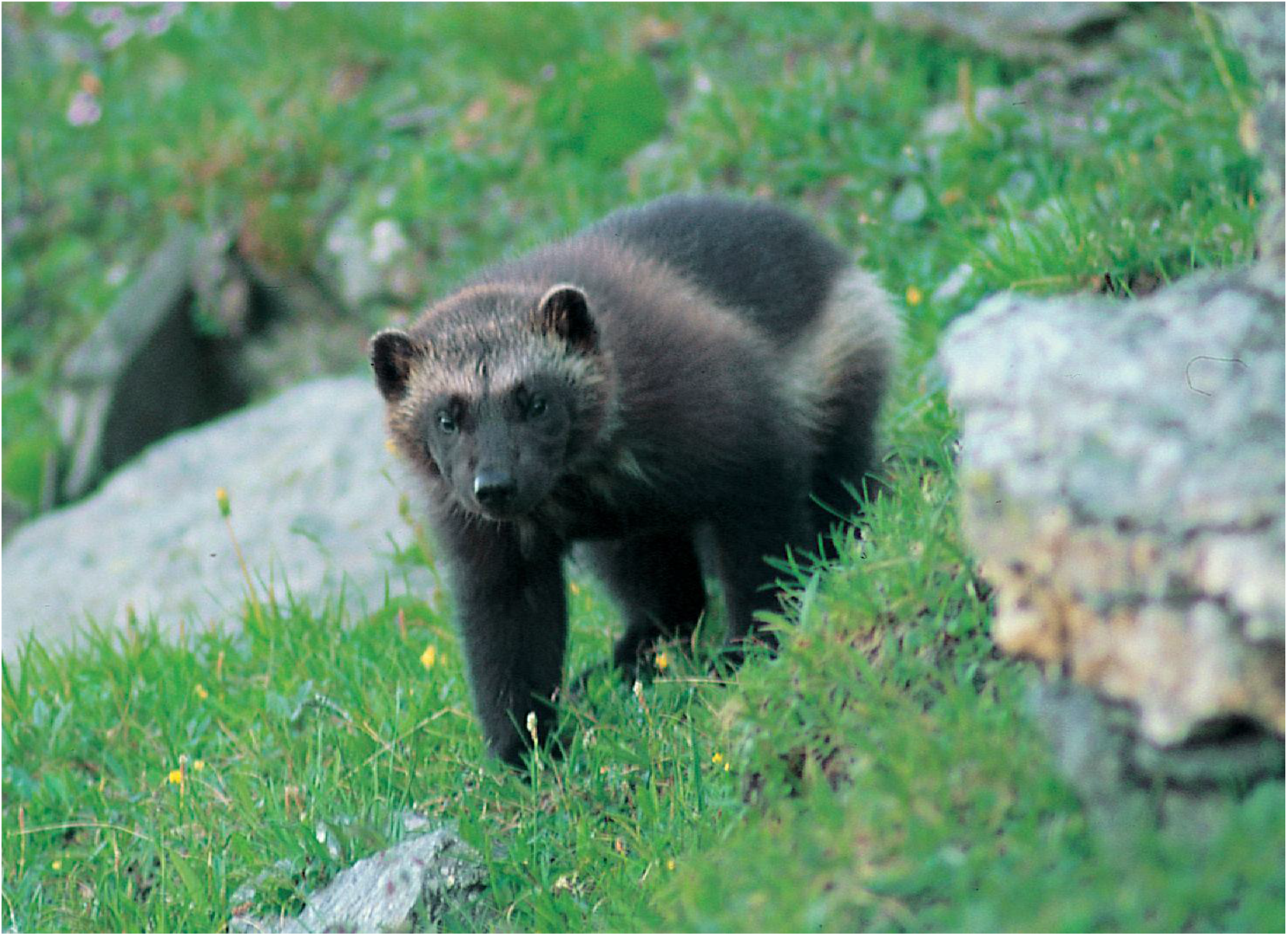
A fennoscandian wolverine (*Gulo gulo gulo*) photographed in Bardu municipality in the summer of 1996. Photo: Roy Andersen.

## Material and Methods

### Sample acquisition and DNA extraction

A tissue sample from a male *G. g. gulo* specimen (RovbaseID: M536555) was collected from Gallonjuonne, Saltdal municipality, Nordland county, Norway (66.90°N, 16.00°E), on 27 May 2025. The specimen was shot by national management authorities as part of damage control. A muscle tissue sample was taken from the heart and preserved in 96% ethanol, stored in a refrigerator for 48h at 4 °C before shipped to the laboratory under EBP-Nor isolate number EBPN_VP_8038_01.

The DNA isolation process for PacBio long-read sequencing started from 60 mg ethanol preserved heart tissue which were split over two BioMasher II tubes and crushed using the Powermasher II tissue disruptor (Nippi Inc.) before proceeding with lysis and DNA isolation with the Nanobind Pan DNA tissue kit (PacBio), following the manufacturer’s recommendations. Quality check of the amount, purity and integrity of the isolated DNA was performed using a combination of Qubit BR DNA quantification assay kit (Thermo Fisher), Nanodrop (Thermo Fisher), and Fragment Analyser (DNA HS 50kb large fragment kit, Agilent Tech.).

### Library preparation and sequencing for *de novo* assembly

Before PacBio HiFi library preparation, DNA was purified an additional time using AMPure PB beads (1:1 ratio). Purified HMW DNA was sheared initially using pipette shearing in 300 µl volume on a Hamilton MLPrep liquid handler (Hamilton Inc) but it resulted in long fragments unsuited for further prepping. Hence, we utilized the Megaruptor3 (Diagenode) with speedcode setting of 32 to generate average fragment sizes of 15-20 kbp large fragments optimal for Pacbio sequencing. A HiFi library was prepared following the PacBio protocol for HiFi library preparation using the SMRTbell® Prep Kit 3.0. The final HiFi library was size-selected with a 3.1x diluted Ampure beads protocol (Pacbio) and sequencing was performed on one SMRT cell (using the associated Revio Binding kit and Sequencing chemistry) on the PacBio Revio instrument (PacBio Inc) located at the Norwegian Sequencing Centre (University of Oslo, Norway).

A formalin cross-linked proximity ligated Hi-C library was prepared by using 30 mg ethanol preserved heart tissue into the Arima High Coverage HiC kit (Arima Genomics), and following the manufacturer’s user guide for low input preparations (Document Part Number A160162 v01), followed by Illumina short read library preparation with the Arima library prep Module and associated user guide (A101030, A303011, A160186 v02). Final library quality was validated using the equipment mentioned above and quantified using a Kapa Library quantification kit for Illumina (Roche Diagnostics) on a LightCyler-96 qPCR instrument (Roche Diagnostics). The library was sequenced with other libraries on an Illumina NovaSeq X (lllumina Inc) with 2*150 bp paired end mode at the Norwegian Sequencing Centre (https://www.sequencing.uio.no).

### Genome assembly and curation, annotation and evaluation

A full list of relevant software tools and versions is presented in Table 1. We assembled the species using a pre-release of the EBP-Nor genome assembly pipeline (https://github.com/ebp-nor/GenomeAssembly). KMC (Kokot et al., 2017) was used to count k-mers of size 32 in the PacBio HiFi reads, excluding k-mers occurring more than 10,000 times. GenomeScope (Ranallo-Benavidez et al., 2020) was run as part of the pipeline on the k-mer histogram output from KMC and was included in the methods for completeness. Ploidy level was investigated using Smudgeplot (Ranallo-Benavidez et al., 2020). HiFiAdapterFilt (Sim et al., 2022) was applied on the HiFi reads to remove possible remnant PacBio adapter sequences. The filtered HiFi reads were assembled using Hifiasm (Cheng et al., 2021) with Hi-C integration resulting in a pair of haplotype-resolved assemblies, pseudo-haplotype one (hap1) and pseudo-haplotype two (hap2). Unique k-mers in each assembly/pseudo-haplotype were identified using meryl (Rhie et al., 2020) and used to create two sets of Hi-C reads, one without any k-mers occurring uniquely in hap1 and the other without k-mers occurring uniquely in hap2.

**Table 1.** Software tools: versions and sources.

| Software tool | Version | Source |
|---|---|---|
| AGAT | 1.5.1 | https://github.com/NBISweden/AGAT |
| BEDTools | 2.31.1 | https://github.com/arq5x/bedtools2 |
| blobtk | 0.5.8 | https://github.com/blobtoolkit/blobtk |
| BlobToolKit | 4.1.7 | https://github.com/blobtoolkit/blobtoolkit |
| BUSCO | 5.8.3 | https://gitlab.com/ezlab/busco |
| BWA-MEM | 0.7.18 | https://github.com/lh3/bwa |
| DIAMOND | 2.1.8 | https://github.com/bbuchfink/diamond |
| EMBLmyGFF3 | 2.2 | https://github.com/NBISweden/EMBLmyGFF3 |
| EvidenceModeler | 1.1.0 | https://github.com/EVidenceModeler/EVidenceModeler |
| FCS-GX | 0.4.0 | https://github.com/ncbi/fcs |
| Funannotate | 1.8.17 | https://github.com/nextgenusfs/funannotate |
| GALBA | 1.0.9 | https://github.com/Gaius-Augustus/GALBA |
| GenomeScope | 2.0 | https://github.com/tbenavi1/genomescope2.0 |
| HiFiAdapterFilt | 2.0.0 | https://github.com/sheinasim/HiFiAdapterFilt |
| hifiasm | 0.25.0 | https://github.com/chhylp123/hifiasm |
| InterProScan | 5.62-94 | https://www.ebi.ac.uk/interpro/search/sequence/ |
| KMC | 3.1.1 | https://github.com/refresh-bio/KMC |
| Merqury | 1.3 | https://github.com/marbl/merqury |
| meryl | 1.3.0 | https://github.com/marbl/meryl |
| minimap2 | 2.28 | https://github.com/lh3/minimap2 |
| miniprot | 0.18 | https://github.com/lh3/miniprot |
| oatk | 1.0 | https://github.com/c-zhou/oatk |
| pafpy | 0.2.0 | https://github.com/mbhall88/pafpy |
| PretextMap | 0.1.9 | https://github.com/wtsi-hpag/PretextMap |
| PretextSnapshot | - | https://github.com/wtsi-hpag/PretextSnapshot |
| PretextView | 0.2.5 | https://github.com/wtsi-hpag/PretextView |
| pyGenomeViz | 1.6.1 | https://github.com/moshi4/pyGenomeViz |
| Rapid curation 2.0 | 964d17e997e00c69f25940cf96d3658bda631147 | https://github.com/Nadolina/Rapid-curation-2.0, |
| RED | 2018.09.10 | https://github.com/BioinformaticsToolsmith/Red |
| samtools | 1.17 | https://github.com/samtools/samtools |
| smudgeplot | 0.2.5 | https://github.com/KamilSJaron/smudgeplot |
| YaHS | 1.2.2 | https://github.com/c-zhou/yahs |

K-mer filtered Hi-C reads were aligned to each scaffolded assembly using BWA-MEM (H. Li, 2013) with `-5SPM` options. The alignments were sorted based on name using samtools (H. Li et al., 2009) before applying samtools fixmate to remove unmapped reads and secondary alignments and to add mate score, and samtools markdup to remove duplicates. The resulting BAM files were used to scaffold the two assemblies using YaHS (Zhou et al., 2023) with default options. FCS-GX (Astashyn et al., 2024) was used to search for putative contamination and contaminated sequences were removed. The mitochondrion was searched for in reads using Oatk (Zhou et al., 2025). The assemblies were manually curated using PretextView and Rapid curation 2.0. Chromosomes were identified by inspecting the Hi-C contact map in PretextView.

We annotated the genome assemblies using a pre-release version of the EBP-Nor genome annotation pipeline (https://github.com/ebp-nor/GenomeAnnotation). First, AGAT (https://zenodo.org/record/7255559) `agat_sp_keep_longest_isoform.pl` and `agat_sp_extract_sequences.pl` were used on the *Homo sapiens* (GRCh38) genome assembly and annotation to generate one protein (the longest isoform) per gene. Miniprot (H. Li, 2023) was used to align the proteins to the curated assemblies. UniProtKB/Swiss-Prot (UniProt Consortium, 2023) release 2025_03 in addition to the Mammalia part of OrthoDB v12 (Tegenfeldt et al., 2025) were also aligned separately to the assemblies. RED (Girgis, 2015) was run via redmask (https://github.com/nextgenusfs/redmask) on the assemblies to mask repetitive areas. In addition to gene models from protein alignments, GALBA (Brůna et al., 2023; Buchfink et al., 2015; Hoff & Stanke, 2019; H. Li, 2023; Stanke et al., 2006) was run with the GRCh38 proteins using the miniprot mode on the masked assemblies to generate *ab initio* predicted gene models. Helixer (Holst et al., 2025) was run using the vertebrate-specific model (vertebrate_v0.3_m_0080).

The `funannotate-runEVM.py` script from Funannotate was used to run EvidenceModeler (Haas et al., 2008) on the alignments of GRCh38 proteins, UniProtKB/Swiss-Prot proteins, Mammalia proteins and the predicted genes from GALBA. The resulting predicted proteins were compared to the protein repeats that Funannotate distributes using DIAMOND blastp and the predicted genes were filtered based on this comparison using AGAT. The filtered proteins were compared to the UniProtKB/Swiss-Prot release 2025_03 using DIAMOND (Buchfink et al., 2015) `blastp` to find gene names and InterProScan (Jones et al., 2014) was used to discover functional domains. AGATs `agat_sp_manage_functional_annotation.pl` was used to attach the gene names and functional annotations to the predicted genes. EMBLmyGFF3 (Norling et al., 2018) was used to combine the FASTA and GFF3 files into a EMBL format for submission to ENA.

All the evaluation tools have also been implemented in a pipeline, similar to assembly and annotation (https://github.com/ebp-nor/GenomeEvaluation). Merqury (Rhie et al., 2020) was used to assess the completeness and quality of the genome assemblies by comparing them to the k-mer content of both the Hi-C reads and PacBio HiFi reads. BUSCO (Manni et al., 2021) was used to assess the completeness of the genome assemblies by comparing against the expected gene content in the eudicots lineage. Gfastats (Formenti et al., 2022) was used to output different assembly statistics of the assemblies. To create snail plots (showing assembly statistics in a condensed plot) and blobplots (showing size, coverage and best taxonomical hit for all sequences in a plot), all sequences in each were compared to UniProt reference proteomes (release 2023_3) (UniProt Consortium, 2025) using mmseqs (Steinegger & Söding, 2017). Further, sequences without hits to reference proteomes were identified, and then compared to the NCBI nt database (downloaded December 2023) (Sayers et al., 2025) using mmseqs. Both the UniProt reference proteomes and NCBI nt databases contain taxonomical information (each entry is associated with a particular species). Together with BUSCO results and coverage (BAM file by mapping PacBio to the assembly with minimap2 (H. Li, 2017)), the hits from the databases were imported into a BlobDir with BlobToolKit and BlobTools2 (Laetsch & Blaxter, 2017). BlobToolKit and BlobTools2 (Laetsch & Blaxter, 2017), in addition to blobtk, were used to create snail plots and blobplots. To generate the Hi-C contact map image, the Hi-C reads were mapped to the assemblies using BWA-MEM (H. Li, 2013) using the same approach as above. Finally, PretextMap was used to create a contact map which was visualized using PretextSnapshot.

To compare the assemblies generated in this study to other available wolverine genome assemblies, we downloaded GCA_024509225.1 and GCA_024510155.1 from NCBI and aligned these to each other and the assemblies generated here using minimap2 (H. Li, 2017) resulting in pairwise PAF files. For visualisation purposes, scaffolds in the downloaded assemblies were ordered according to hap1, and only sequences longer than 5 Mb were included in plots, or only chromosomes if that information was available. PAF files were processed using pafpy and pyGenomeViz was used for plotting (see Zenodo repository listed under Data availability for the script used in plotting).

The text in Methods and parts of Results is based on a template we use for all the species we publish in the EBP-Nor project.

## Results

### *De novo* genome assembly and annotation

The genome of *Gulo gulo gulo* was assembled from a total of 40-fold coverage in Pacific Biosciences single-molecule HiFi long reads and 99-fold coverage in Arima Hi-C reads resulting in two haplotype-separated assemblies. The estimated genome size was 2,270 Mb (Supplementary Figure 1). The final assemblies have total lengths of 2,523.2 Mb and 2,288.4 Mb (Table 2 and Figure 2), respectively. Pseudo-haplotypes one (hap1) and two (hap2) have scaffold N50 size of 163.6 Mb and 162.6 Mb, respectively, and contig N50 of 82.2 Mb and 81.4 Mb, respectively (Table 2 and Fig. 2). Twenty pseudo-chromosomes were identified in both pseudo-haplotypes, in addition to X and Y in hap1. Chromosomes are named according to size.

**Table 2:** Genome data for *Gulo gulo gulo*.

| **Project accession data** |  |  |
|---|---|---|
| Species | *Gulo gulo gulo* |  |
| Specimen | mGulGul1 |  |
| NCBI Taxonomy ID | 2888764 |  |
| BioProject | PRJEB125773 |  |
| BioSample ID | SAMEA123427954 |  |
| Isolate information | Male, muscle |  |
| **Raw data accessions** |  |  |
| PacBio HiFi reads | ERR17844416 and ERR17845569 | 2 PACBIO_SMRT (Revio) run: 6.2 M reads, 101.1 Gb |
| Hi-C Illumina reads | ERR17844179 and ERR17844182 | 2 ILLUMINA (Illumina NovaSeq X run: 789 M pairs of reads, 238.3 Gb |
| **Genome assembly metrics** |  |  |
| HiFi read coverage | 40x |  |
| Assembly accession | GCA_988242245 | GCA_988242235 |
| Assembly identifier | mGulGul1.1.hap1 | mGulGul1.1.hap2 |
| Span (Mb) | 2,523.2 | 2,288.4 |
| Number of contigs | 630 | 420 |
| Contig N50 length (Mb) | 82.2 | 81.4 |
| Longest contig (Mb) | 163.6 | 162.6 |
| Number of gaps | 67 | 130 |
| Number of scaffolds | 563 | 293 |
| Scaffold N50 length (Mb) | 140.0 | 141.9 |
| Longest scaffold (Mb) | 222.0 | 213.2 |
| BUSCO* (genome) | C:98.2%[S:97.2%,D:1.0%],F:1.2%,M:0.6 | C:88.9%[S:88.2%,D:0.7%],F:1.1%,M:10.0% |
| Consensus quality (QV) compared to Hi-C (compared to HiFi) | 50.0 (68.6) | 51.0 (70.8) |
| Both assemblies | 50.5 (69.5) |  |
| *k*-mer completeness (%) compared to Hi-C (compared to HiFi) | 97.4 (98.3) | 90.9 (91.4) |
| Both assemblies | 98.7 (99.7) |  |
| Percentage of assembly mapped to chromosomes | 95.8 | 97.8 |
| Sex chromosomes | X and Y |  |
| Organelles | MT |  |
| **Genome annotation metrics** |  |  |
| Number of protein-coding genes | 20,337 | 17,692 |
| Number of protein-coding genes with functional domain** | 20,018 | 17,453 |
| Number of protein-coding genes with gene names | 19,248 | 16,846 |
| BUSCO* | C:96.8%[S:96.0%,D:0.8%],F:0.5%,M:2.7%, | C:87.2%[S:86.7%,D:0.5%],F:0.5%,M:12.2% |

\* BUSCO scores based on the carnivora BUSCO set using v6.0.0. C = complete [S = single copy, D = duplicated], F = fragmented, M = missing. \*\*Number of genes annotated with a functional domain as found by InterProScan.

**Figure 2:**
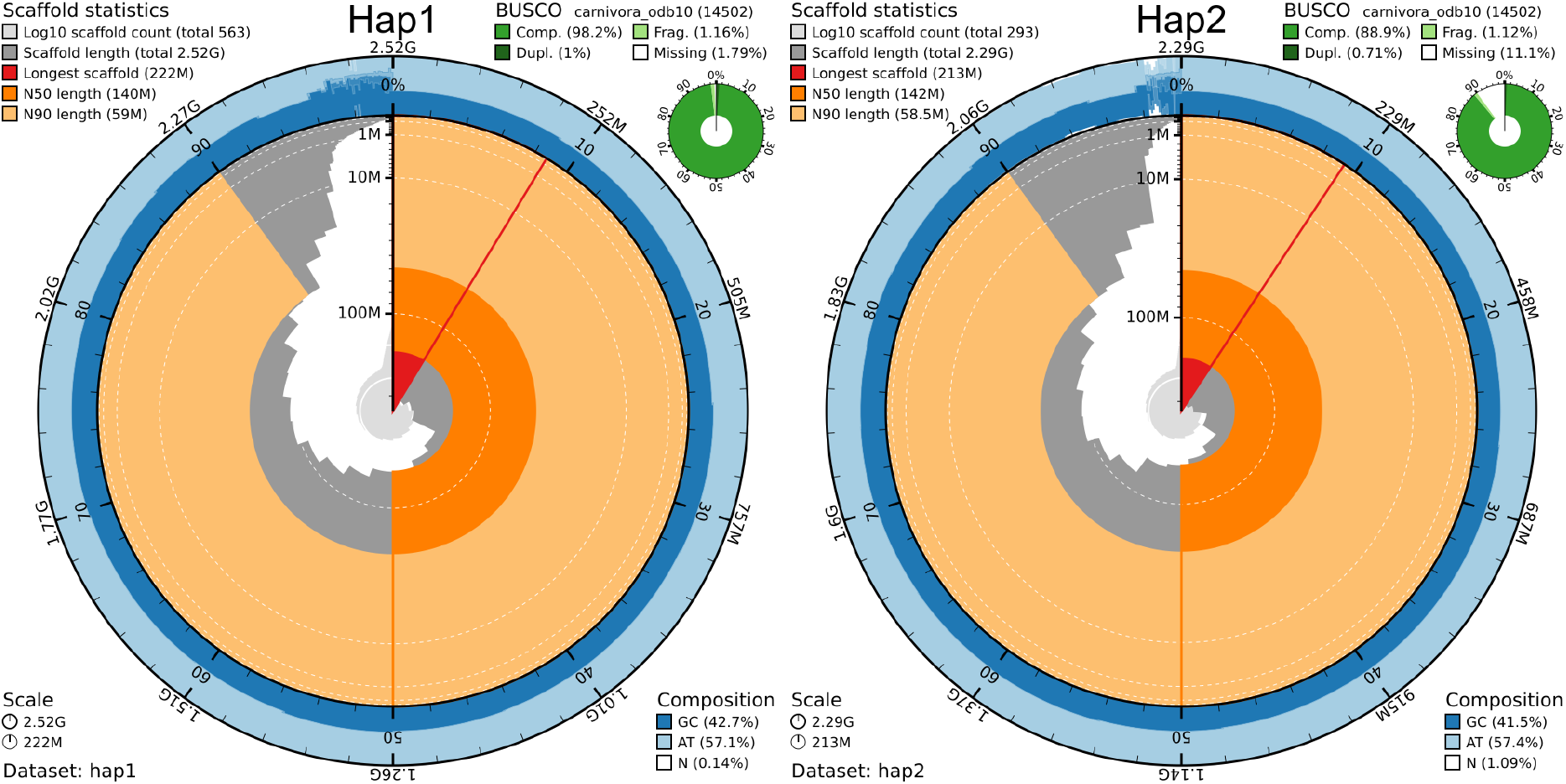
Metrics of the genome assemblies of *Gulo gulo gulo* hap1 and hap2. A) Hap1. B) Hap2. The BlobToolKit Snailplots show N50 metrics and BUSCO gene completeness. The two outermost bands of the circle signify GC versus AT composition at 0.1% intervals. Light orange shows the N90 scaffold length, while the deeper orange is N50 scaffold length. The red line shows the size of the largest scaffold. All the scaffolds are arranged in a clockwise manner from the largest to the smallest and are shown in darker gray with white lines at different orders of magnitude, while the light gray shows cumulative count of scaffolds.

When compared to a k-mer database of the Hi-C reads, hap1 had a k-mer completeness of 97.4%, hap2 of 90.9%, and combined they have a completeness of 98.7%. Further, hap1 had an assembly consensus quality value (QV) of 50.0 and hap2 of 51.0, where a QV of 50 corresponds to one error every 100,000 bp, or 99.999% accuracy compared to a k-mer database of the Hi-C reads (QV 68.6 and 70.8, respectively, compared to a k-mer database of the HiFi reads) (Table 2). Plot of k-mers in assemblies compared to k-mers in reads shows that k-mers occurring one or a few times in reads (likely errors) do not occur in the assemblies (Supplementary Figure 3).

A total of 20,337 and 17,692 protein-coding genes were annotated in hap1 and hap2, respectively (Table 2).

The Hi-C contact maps show clear separation of chromosomes into homologous sets (Supplementary Figure 2). While the blobplots (Supplementary Figure 4) indicate some possible contamination, this is due to incomplete reference databases and how the plots were created (Supplementary Text 1).

To further assess the quality of the assemblies generated here, we plotted them against other chromosome-level genome assemblies available for wolverine, GCA_024509225.1 and GCA_024510155.1 (both from Lok et al. (2022); Figure 3). There are two other genome assemblies made public, GCA_90000637Lok et al. (2022)5.2 and GCA_023973185.1, but these are based on Illumina reads without long range scaffolding information and are therefore much more fragmented with scaffolds in the tens of thousands (Table 3). These were not included.

**Figure 3:**
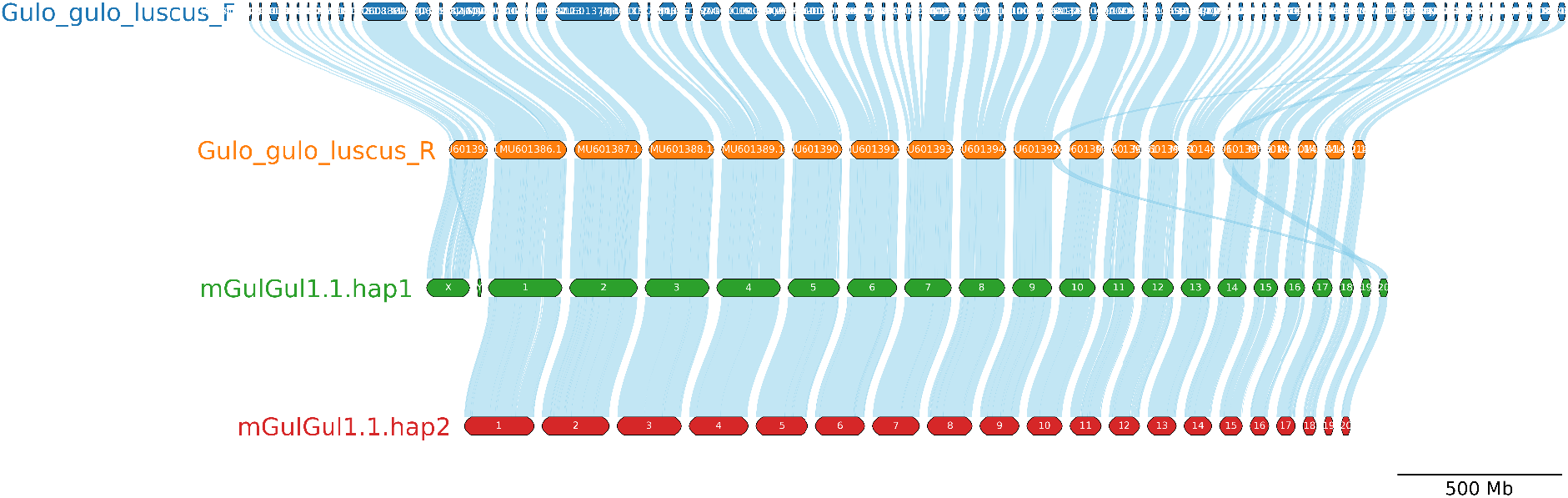
Synteny plot of available wolverine genome assemblies. The pseudo-haplotypes (mGulGul1.1.hap1 and mGulGul1.1.hap2) have been scaffolded with Hi-C data, while Gulo_gulo_luscus_R is a version of Gulo_gulo_luscus_F (Lok et al. 2022) that has been scaffolded based on alignments to ermine (*Mustela erminea*), Eurasian river otter (*Lutra lutra*) and the dingo (*Canis lupus dingo)*.

**Table 3:** Genome statistics for available wolverine genome assemblies.

| Assembly | Accessions | Length (Mb) | N50 scaffold (Mb) | N50 contig (Mb) |
|---|---|---|---|---|
| mGulGul1.1.hap1 | GCA_988242245.1 | 2,523 | 140.0 | 82.2 |
| mGulGul1.1.hap2 | GCA_988242235.1 | 2,288 | 141.9 | 81.4 |
| Gulo_gulo_luscus_R | GCA_024510155.1 | 2,388 | 144.0 | 36.6 |
| Gulo_gulo_luscus_F | GCA_024509225.1 | 2,388 | 50.8 | 36.6 |
| Gulo_gulo_luscus_A | GCA_023973185.1 | 2,249 | 0.05 | 0.02 |
| Gulo_2.2 | GCA_900006375.2 | 2,423 | 0.178 | 0.004 |

## Discussion

The genome assembly presented here represents a chromosome level and pseudo-haplotype-resolved assembly more complete than any wolverine genome assemblies generated so far and with both the X and Y sex chromosomes resolved (Table 3). A previous project generated a chromosome-level assembly based on the North American subspecies, *Gulo gulo luscus* (Lok et al., 2022). They used alignments to ermine (*Mustela erminea*), Eurasian river otter (*Lutra lutra*) and the dingo (*Canis lupus dingo*) as reference-based scaffolding (Gulo_gulo_luscus_R). This resulted in a genome assembly with 18 autosomal chromosomes plus X without haplotype separation (Figure 3). The number of chromosomes in wolverine is 42 (21 chromosome pairs). Compared to the assemblies generated here, the Gulo_gulo_luscus_R assembly has the two smallest chromosomes fused to two different larger chromosomes (Figure 3). This is likely to be the result of using species with different chromosome numbers and therefore different architecture, as references with 44, 36 and 78 for ermine, eurasian river otter and dingo, respectively (Lok et al., 2022). The assemblies generated here contain the correct number of chromosomes, as well as a separate Y chromosome not fused with the autosomes.

The wolverine genome is the first chromosome-level assembly of the Eurasian subspecies (*G. gulo gulo*). It, and more complete genome assemblies of relatives, will give further insights into the evolution of Mustelidae (Derežanin et al., 2022). Further, it will be a valuable reference for population studies and conservation efforts.

## Acknowledgments

This project received data management and infrastructure support from ELIXIR Norway, supported by the Research Council of Norway’s grants 322392 and 350529, the University of Bergen, the University of Oslo, the Arctic University of Norway in Tromsø, the Norwegian University of Science and Technology, and the Norwegian University of Life Sciences: NMBU. The authors acknowledge support from the National Infrastructure for High Performance Computing and resources provided by Sigma2 as well as Data Storage in Norway (project NN8013K) for computational work. The Norwegian Sequencing Centre generated the sequencing data used in this project (http://sequencing.uio.no). We thank the Norwegian Nature Inspectorate (SNO) for sample acquisition.

## Data availability

Data generated for this study are available under ENA BioProject PRJEB125773. PacBio sequencing data for the Eurasian wolverine (ENA BioSample: SAMEA123427954) are deposited in ENA under ERR17844416 and ERR17845569, whereas Illumina Hi-C sequencing data are deposited in ENA under ERR17844179 and ERR17844182. Pseudo-haplotype one can be found in ENA at PRJEB125603, whereas hap2 is PRJEB125604.

The gene annotations (and script for plotting synteny) are available at https://doi.org/10.5281/zenodo.23262416.

## References

Abramov, A. (2015, May 15). IUCN Red List of Threatened Species: Gulo gulo. https://www.iucnredlist.org/en

Andrén, H., Persson, J., Mattisson, J., & Danell, A. C. (2011). Modelling the combined effect of an obligate predator and a facultative predator on a common prey: lynx Lynx lynx and wolverine Gulo gulo predation on reindeer Rangifer tarandus. Wildlife Biology, 17(1), 33–43. 10.2981/10-065

Aronsson, M., & Persson, J. (2018). Female breeding dispersal in wolverines, a solitary carnivore with high territorial fidelity. European Journal of Wildlife Research, 64(1), 7. 10.1007/s10344-018-1164-3

Astashyn, A., Tvedte, E. S., Sweeney, D., Sapojnikov, V., Bouk, N., Joukov, V., Mozes, E., Strope, P. K., Sylla, P. M., Wagner, L., Bidwell, S. L., Brown, L. C., Clark, K., Davis, E. W., Smith-White, B., Hlavina, W., Pruitt, K. D., Schneider, V. A., & Murphy, T. D. (2024). Rapid and sensitive detection of genome contamination at scale with FCS-GX. Genome Biology, 25(1), 60. 10.1186/s13059-024-03198-7

Aubry, K. B., McKelvey, K. S., & Copeland, J. P. (2007). Distribution and broadscale habitat relations of the wolverine in the contiguous United States. The Journal of Wildlife Management, 71(7), 2147–2158. 10.2193/2006-548

Aubry, K. B., Raley, C. M., Shirk, A. J., McKelvey, K. S., & Copeland, J. P. (2023). Climatic conditions limit wolverine distribution in the Cascade Range of southwestern North America. Canadian Journal of Zoology, 101(2), 95–113. 10.1139/cjz-2022-0005

Brůna, T., Li, H., Guhlin, J., Honsel, D., Herbold, S., Stanke, M., Nenasheva, N., Ebel, M., Gabriel, L., & Hoff, K. J. (2023). Galba: genome annotation with miniprot and AUGUSTUS. BMC Bioinformatics, 24(1), 327. 10.1186/s12859-023-05449-z

Buchfink, B., Xie, C., & Huson, D. H. (2015). Fast and sensitive protein alignment using DIAMOND. Nature Methods, 12(1), 59–60. 10.1038/nmeth.3176

Cheng, H., Concepcion, G. T., Feng, X., Zhang, H., & Li, H. (2021). Haplotype-resolved de novo assembly using phased assembly graphs with hifiasm. Nature Methods, 18(2), 170–175. 10.1038/s41592-020-01056-5

Copeland, J. P., McKelvey, K. S., Aubry, K. B., Landa, A., Persson, J., Inman, R. M., Krebs, J., Lofroth, E., Golden, H., Squires, J. R., Magoun, A., Schwartz, M. K., Wilmot, J., Copeland, C. L., Yates, R., Kojola, I., & May, R. (2010). The bioclimatic envelope of the wolverine (Gulo gulo): do climatic constraints limit its geographic distribution? Canadian Journal of Zoology. 88: 233–246., 233–246. https://research.fs.usda.gov/treesearch/34712

Copeland, J. P., & Whitman, J. S. (2003). Wolverine (Gulo gulo). In G. A. Feldhamer, B. C. Thompson, & J. A. Chapman (Eds.), Wild Mammals of North America: Biology, Management, and Economics (pp. 672–682). Johns Hopkins University Press/The Inter-Ame.

DereŽanin, L., BlaŽytė, A., Dobrynin, P., Duchêne, D. A., Grau, J. H., Jeon, S., Kliver, S., Koepfli, K.-P., Meneghini, D., Preick, M., Tomarovsky, A., Totikov, A., Fickel, J., & Förster, D. W. (2022). Multiple types of genomic variation contribute to adaptive traits in the mustelid subfamily Guloninae. Molecular Ecology, 31(10), 2898–2919. 10.1111/mec.16443

Ekblom, R., Brechlin, B., Persson, J., Smeds, L., Johansson, M., Magnusson, J., Flagstad, Ø., & Ellegren, H. (2018). Genome sequencing and conservation genomics in the Scandinavian wolverine population: Conservation Genomics. Conservation Biology: The Journal of the Society for Conservation Biology, 32(6), 1301–1312. 10.1111/cobi.13157

Fisher, J. T., Murray, S., Barrueto, M., Carroll, K., Clevenger, A. P., Hausleitner, D., Harrower, W., Heim, N., Heinemeyer, K., Jacob, A. L., Jung, T. S., Kortello, A., Ladle, A., Long, R., MacKay, P., & Sawaya, M. A. (2022). Wolverines (Gulo gulo) in a changing landscape and warming climate: A decadal synthesis of global conservation ecology research. Global Ecology and Conservation, 34(e02019), e02019. 10.1016/j.gecco.2022.e02019

Formenti, G., Abueg, L., Brajuka, A., Brajuka, N., Gallardo-Alba, C., Giani, A., Fedrigo, O., & Jarvis, E. D. (2022). Gfastats: conversion, evaluation and manipulation of genome sequences using assembly graphs. Bioinformatics (Oxford, England), 38(17), 4214–4216. 10.1093/bioinformatics/btac460

Girgis, H. Z. (2015). Red: an intelligent, rapid, accurate tool for detecting repeats de-novo on the genomic scale. BMC Bioinformatics, 16(1), 227. 10.1186/s12859-015-0654-5

Haas, B. J., Salzberg, S. L., Zhu, W., Pertea, M., Allen, J. E., Orvis, J., White, O., Buell, C. R., & Wortman, J. R. (2008). Automated eukaryotic gene structure annotation using EVidenceModeler and the Program to Assemble Spliced Alignments. Genome Biology, 9(1), R7. 10.1186/gb-2008-9-1-r7

Hoff, K. J., & Stanke, M. (2019). Predicting genes in single genomes with AUGUSTUS. Current Protocols in Bioinformatics, 65(1), e57. 10.1002/cpbi.57

Hohenlohe, P. A., Funk, W. C., & Rajora, O. P. (2021). Population genomics for wildlife conservation and management. Molecular Ecology, 30(1), 62–82. 10.1111/mec.15720

Holst, F., Bolger, A. M., Kindel, F., Günther, C., Maß, J., Triesch, S., Kiel, N., Saadat, N., Ebenhöh, O., Usadel, B., Schwacke, R., Weber, A. P. M., Bolger, M. E., & Denton, A. K. (2025). Helixer: ab initio prediction of primary eukaryotic gene models combining deep learning and a hidden Markov model. Nature Methods. 10.1038/s41592-025-02939-1

Hyvärinen, E., Juslén, A., Kemppainen, E., Uddström, A., & Liukko, U.-M. (Eds.). (2019). Suomen lajien uhanalaisuus : Punainen kirja 2019 (The 2019 Red List of Finnish Specie). Ympäristöministeriö & Suomen ympäristökeskus. https://helda.helsinki.fi/items/2ec69a48-d943-488c-927f-19bbf9f92cb5

Jones, P., Binns, D., Chang, H.-Y., Fraser, M., Li, W., McAnulla, C., McWilliam, H., Maslen, J., Mitchell, A., Nuka, G., Pesseat, S., Quinn, A. F., Sangrador-Vegas, A., Scheremetjew, M., Yong, S.-Y., Lopez, R., & Hunter, S. (2014). InterProScan 5: genome-scale protein function classification. Bioinformatics (Oxford, England), 30(9), 1236–1240. 10.1093/bioinformatics/btu031

Kokot, M., Dlugosz, M., & Deorowicz, S. (2017). KMC 3: counting and manipulating k-mer statistics. Bioinformatics (Oxford, England), 33(17), 2759–2761. 10.1093/bioinformatics/btx304

Krejsa, D. M., Talbot, S. L., Sage, G. K., Sonsthagen, S. A., Jung, T. S., Magoun, A. J., & Cook, J. A. (2021). Dynamic landscapes in northwestern North America structured populations of wolverines (Gulo gulo). Journal of Mammalogy, 102(3), 891–908. 10.1093/jmammal/gyab045

Laetsch, D. R., & Blaxter, M. L. (2017). BlobTools: Interrogation of genome assemblies. F1000Research, 6, 1287. 10.12688/f1000research.12232.1

Landa, A., Gudvangen, K., Swenson, J. E., & Roskaft, E. (1999). Factors associated with wolverine Gulo gulo predation on domestic sheep. The Journal of Applied Ecology, 36(6), 963–973. 10.1046/j.1365-2664.1999.00451.x

Li, B., Wolsan, M., Wu, D., Zhang, W., Xu, Y., & Zeng, Z. (2014). Mitochondrial genomes reveal the pattern and timing of marten (Martes), wolverine (Gulo), and fisher (Pekania) diversification. Molecular Phylogenetics and Evolution, 80, 156–164. 10.1016/j.ympev.2014.08.002

Li, H. (2013). Aligning sequence reads, clone sequences and assembly contigs with BWA-MEM. In arXiv [q-bio.GN]. arXiv. http://arxiv.org/abs/1303.3997

Li, H. (2017). Minimap2: pairwise alignment for nucleotide sequences. Bioinformatics, 34, 3094–3100. 10.1093/bioinformatics/bty191

Li, H. (2023). Protein-to-genome alignment with miniprot. Bioinformatics (Oxford, England), 39(1), btad014. 10.1093/bioinformatics/btad014

Li, H., Handsaker, B., Wysoker, A., Fennell, T., Ruan, J., Homer, N., Marth, G., Abecasis, G., Durbin, R., & 1000 Genome Project Data Processing Subgroup. (2009). The Sequence Alignment/Map format and SAMtools. Bioinformatics (Oxford, England), 25(16), 2078–2079. 10.1093/bioinformatics/btp352

Lok, S., Lau, T. N. H., Trost, B., Tong, A. H. Y., Wintle, R. F., Engstrom, M. D., Stacy, E., Waits, L. P., Scrafford, M., & Scherer, S. W. (2022). Chromosomal-level reference genome assembly of the North American wolverine (Gulo gulo luscus): a resource for conservation genomics. G3 (Bethesda, Md.), 12(8), jkac138. 10.1093/g3journal/jkac138

Manni, M., Berkeley, M. R., Seppey, M., Simão, F. A., & Zdobnov, E. M. (2021). BUSCO update: Novel and streamlined workflows along with broader and deeper phylogenetic coverage for scoring of eukaryotic, prokaryotic, and viral genomes. Molecular Biology and Evolution, 38(10), 4647–4654. 10.1093/molbev/msab199

Mattisson, J., Andrén, H., Persson, J., & Segerström, P. (2011). Influence of intraguild interactions on resource use by wolverines and Eurasian lynx. Journal of Mammalogy, 92(6), 1321–1330. 10.1644/11-mamm-a-099.1

McKelvey, K. S., Copeland, J. P., Schwartz, M. K., Littell, J. S., Aubry, K. B., Squires, J. R., Parks, S. A., Elsner, M. M., & Mauger, G. S. (2011). Climate change predicted to shift wolverine distributions, connectivity, and dispersal corridors. Ecological Applications, 21(8), 2882–2897. 10.1890/10-2206.1

Norling, M., Jareborg, N., & Dainat, J. (2018). EMBLmyGFF3: a converter facilitating genome annotation submission to European Nucleotide Archive. BMC Research Notes, 11(1), 584. 10.1186/s13104-018-3686-x

Norwegian Biodiversity Information Centre. (2021, November 24). Norwegian Red List for Species 2021. Artsdatabanken. https://lister.artsdatabanken.no/rodlisteforarter/2021/

Persson, J., Ordiz, A., Ladle, A., Andrén, H., & Aronsson, M. (2023). Recolonization following past persecution questions the importance of persistent snow cover as a range limiting factor for wolverines. Global Change Biology, 29(20), 5802–5815. 10.1111/gcb.16908

Ranallo-Benavidez, T. R., Jaron, K. S., & Schatz, M. C. (2020). GenomeScope 2.0 and Smudgeplot for reference-free profiling of polyploid genomes. Nature Communications, 11(1), 1432. 10.1038/s41467-020-14998-3

Rhie, A., Walenz, B. P., Koren, S., & Phillippy, A. M. (2020). Merqury: reference-free quality, completeness, and phasing assessment for genome assemblies. Genome Biology, 21(1), 245. 10.1186/s13059-020-02134-9

Ruggiero, L. F., Mckelvey, K. S., Aubry, K. B., Copeland, J. P., Pletscher, D. H., & Hornocker, M. G. (2007). Wolverine conservation and management. The Journal of Wildlife Management, 71(7), 2145–2146. 10.2193/2007-217

Sayers, E. W., Beck, J., Bolton, E. E., Brister, J. R., Chan, J., Connor, R., Feldgarden, M., Fine, A. M., Funk, K., Hoffman, J., Kannan, S., Kelly, C., Klimke, W., Kim, S., Lathrop, S., Marchler-Bauer, A., Murphy, T. D., O’Sullivan, C., Schmieder, E., … Pruitt, K. D. (2025). Database resources of the National Center for Biotechnology Information in 2025. Nucleic Acids Research, 53(D1), D20–D29. 10.1093/nar/gkae979

Shafer, A. B. A., Wolf, J. B. W., Alves, P. C., Bergström, L., Bruford, M. W., Brännström, I., Colling, G., Dalén, L., De Meester, L., Ekblom, R., Fawcett, K. D., Fior, S., Hajibabaei, M., Hill, J. A., Hoezel, A. R., Höglund, J., Jensen, E. L., Krause, J., Kristensen, T. N., … Zieliński, P. (2015). Genomics and the challenging translation into conservation practice. Trends in Ecology & Evolution, 30(2), 78–87. 10.1016/j.tree.2014.11.009

Sim, S. B., Corpuz, R. L., Simmonds, T. J., & Geib, S. M. (2022). HiFiAdapterFilt, a memory efficient read processing pipeline, prevents occurrence of adapter sequence in PacBio HiFi reads and their negative impacts on genome assembly. BMC Genomics, 23(1), 157. 10.1186/s12864-022-08375-1

Stanke, M., Schöffmann, O., Morgenstern, B., & Waack, S. (2006). Gene prediction in eukaryotes with a generalized hidden Markov model that uses hints from external sources. BMC Bioinformatics, 7(1), 62. 10.1186/1471-2105-7-62

Steinegger, M., & Söding, J. (2017). MMseqs2 enables sensitive protein sequence searching for the analysis of massive data sets. Nature Biotechnology, 35(11), 1026–1028. 10.1038/nbt.3988

Swedish Species Information Centre, Swedish University of Agricultural Sciences. (2025). The Swedish Red List 2020 [Dataset]. Swedish University of Agricultural Sciences. 10.5878/0KX8-XR80

Tegenfeldt, F., Kuznetsov, D., Manni, M., Berkeley, M., Zdobnov, E. M., & Kriventseva, E. V. (2025). OrthoDB and BUSCO update: annotation of orthologs with wider sampling of genomes. Nucleic Acids Research, 53(D1), D516–D522. 10.1093/nar/gkae987

Thiel, A., Evans, A. L., Fuchs, B., Arnemo, J. M., Aronsson, M., & Persson, J. (2019). Effects of reproduction and environmental factors on body temperature and activity patterns of wolverines. Frontiers in Zoology, 16(1), 21. 10.1186/s12983-019-0319-8

UniProt Consortium. (2023). UniProt: The universal protein knowledgebase in 2023. Nucleic Acids Research, 51(D1), D523–D531. 10.1093/nar/gkac1052

UniProt Consortium. (2025). UniProt: The universal protein knowledgebase in 2025. Nucleic Acids Research, 53(D1), D609–D617. 10.1093/nar/gkae1010

van Dijk, J., Andersen, T., May, R., Andersen, R., Andersen, R., & Landa, A. (2008). Foraging strategies of wolverines within a predator guild. Canadian Journal of Zoology, 86(9), 966–975. 10.1139/z08-073

Zhou, C., Brown, M., Blaxter, M., Darwin Tree of Life Project Consortium, McCarthy, S. A., & Durbin, R. (2025). Oatk: a de novo assembly tool for complex plant organelle genomes. Genome Biology, 26(1), 235. 10.1186/s13059-025-03676-6

Zhou, C., McCarthy, S. A., & Durbin, R. (2023). YaHS: yet another Hi-C scaffolding tool. Bioinformatics (Oxford, England), 39(1), btac808. 10.1093/bioinformatics/btac808

Zigouris, J., Schaefer, J. A., Fortin, C., & Kyle, C. J. (2013). Phylogeography and post-glacial recolonization in wolverines (Gulo gulo) from across their circumpolar distribution. PloS One, 8(12), e83837. 10.1371/journal.pone.0083837

