## Supplementary information for "A chromosome-level genome assembly of an endangered large carnivore, the wolverine, *Gulo gulo gulo* (Linnaeus, 1758)"

Authors:
**Ole K. Tørresen [1], Øystein Flagstad [2], Oddmund Kleven [2], Marius A. Strand [1], Morten Skage [1], Giada Ferrari [1], Ave Tooming-Klunderud [1], Kjetill S. Jakobsen [1*]**
Addresses:
1) Department of Biosciences, University of Oslo, Oslo, Norway
2) Norwegian Institute for Nature Research, Trondheim, Norway

### Supplementary

**Supplementary Text 1. Putative contaminants in the genome assemblies.** Supplementary Figure 4 shows that, based on searches in UniProt reference proteomes and NCBI nt, there might be some contamination in the assemblies. When trying to validate these results by isolating sequences with hits towards Nematoda and using the NCBI BLAST website, these sequences have hits against rRNA from Mustelidae species. As mentioned in Material and Methods, the process to generate blobplots involves matching against UniProt reference proteomes first, and the against NCBI nt if a sequence does not have a good hit against a reference proteome. Here, the rRNA sequences from *Gulo gulo* happen to have good matches against Nematoda sequences from UniProt reference proteomes. This is an automated process that incidentally gives wrong results, but manually adjusting this would partly invalidate the process. These databases (NCBI nt and UniProt reference proteomes) do contain errors, and/or, parts of genomes from quite different lineages happen to be quite similar in sequence.

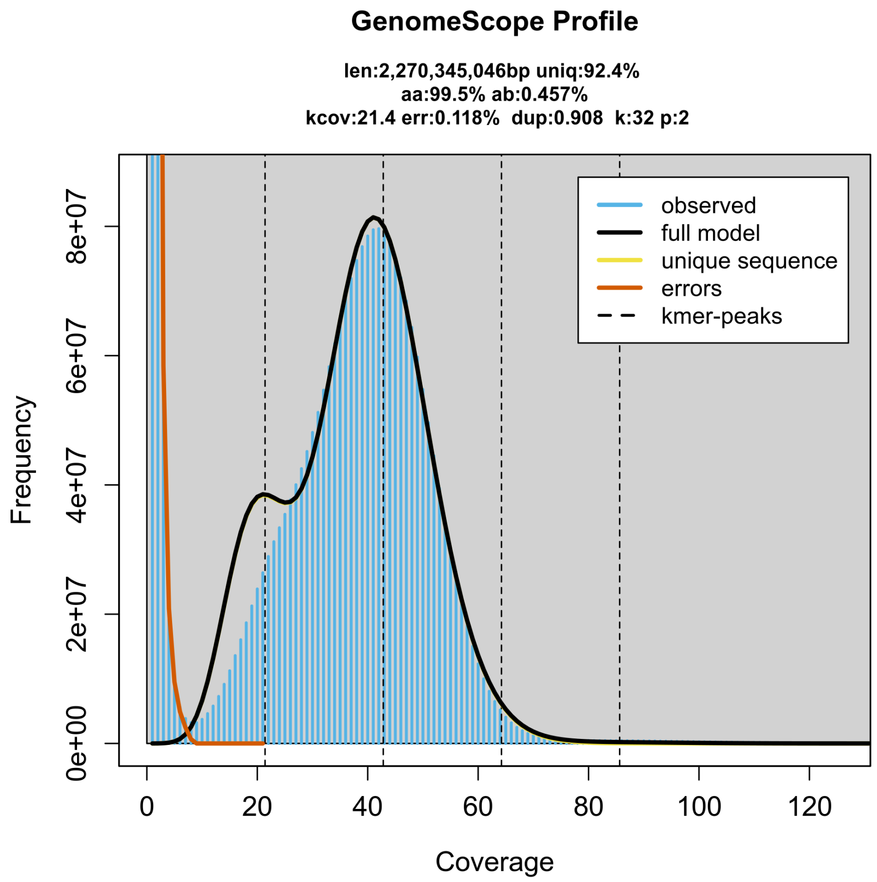


**Supplementary Figure 1. Genome profile of *Gulo gulo gulo.*** GenomeScope profile of the HiFi reads from the sequenced individual. This analysis estimates a 2,270 Mb genome, with 0.457% heterozygosity. The left-hand shoulder of k-mers corresponds to k-mers from heterozygous regions of the genome, while the mean peak is from homozygous regions.

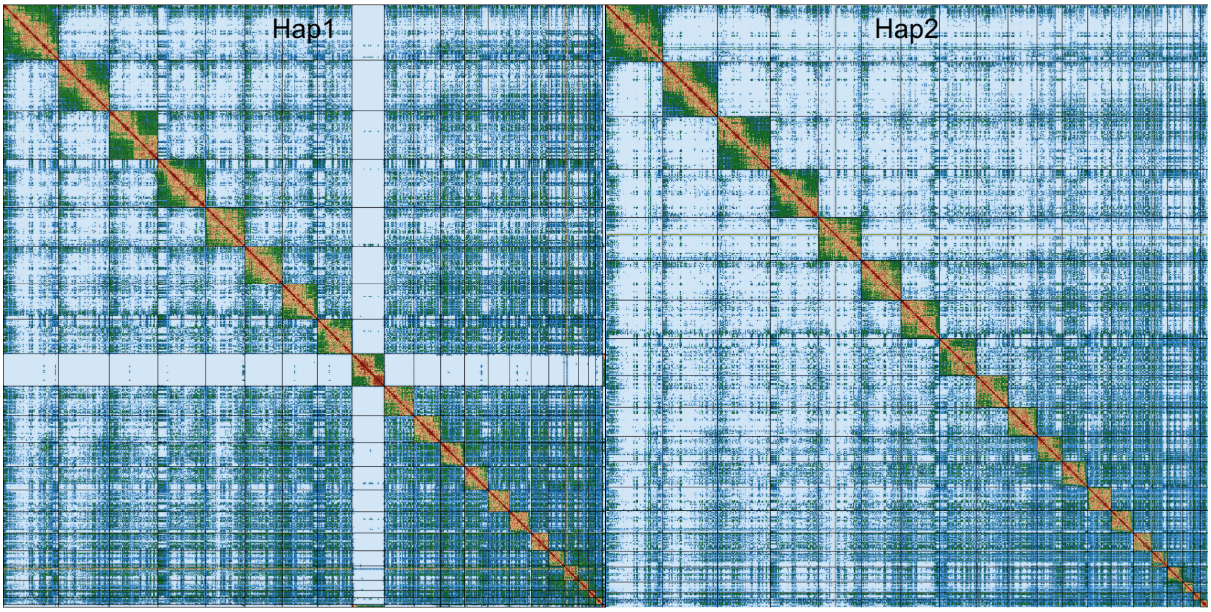


**Supplementary Figure 2: Hi-C contact maps for *Gulo gulo gulo* hap1 and hap2.** The contact map displays interaction frequencies between genomic regions, where darker shades represent a higher number of Hi-C contacts. The axes correspond to the coordinates along each assembly. The Hi-C contact maps were generated by mapping the Hi-C reads to the haplotype genomes using BWA-mem, generating a contact map using PretextMap and visualized using PretextSnapshot.

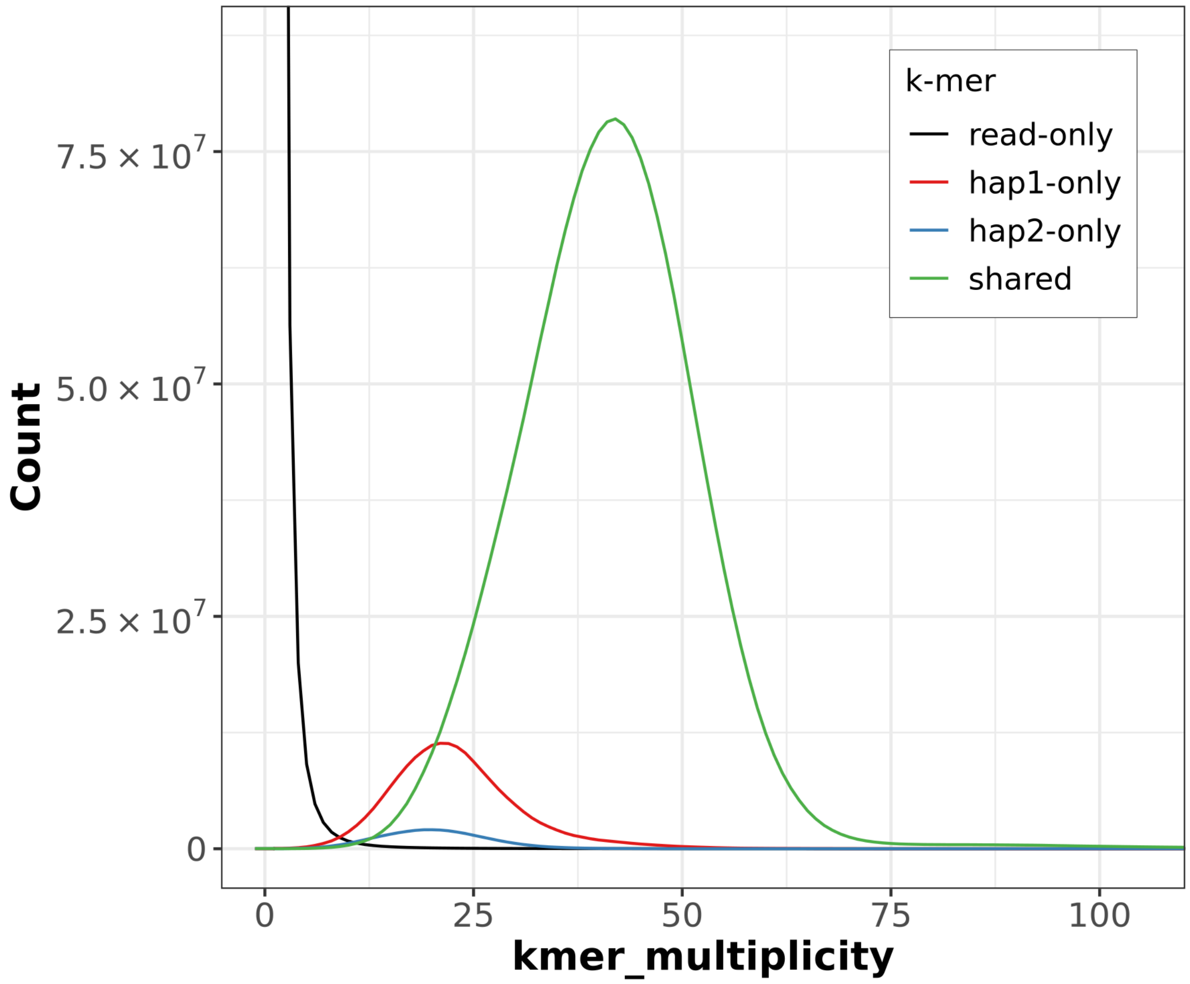


**Supplementary Figure 3: *K*-mer spectra of HiFi reads for *Gulo gulo gulo*.** Distributions of *k*-mers found only in the reads (black), only in haplotype 1 (red), only in haplotype 2 (blue), or in both haplotypes (green). The x-axis represents the number of unique *k*-mers, while the y-axis represents the *k*-mer multiplicity (how often the *k*-mer is found in the set of HiFi reads).

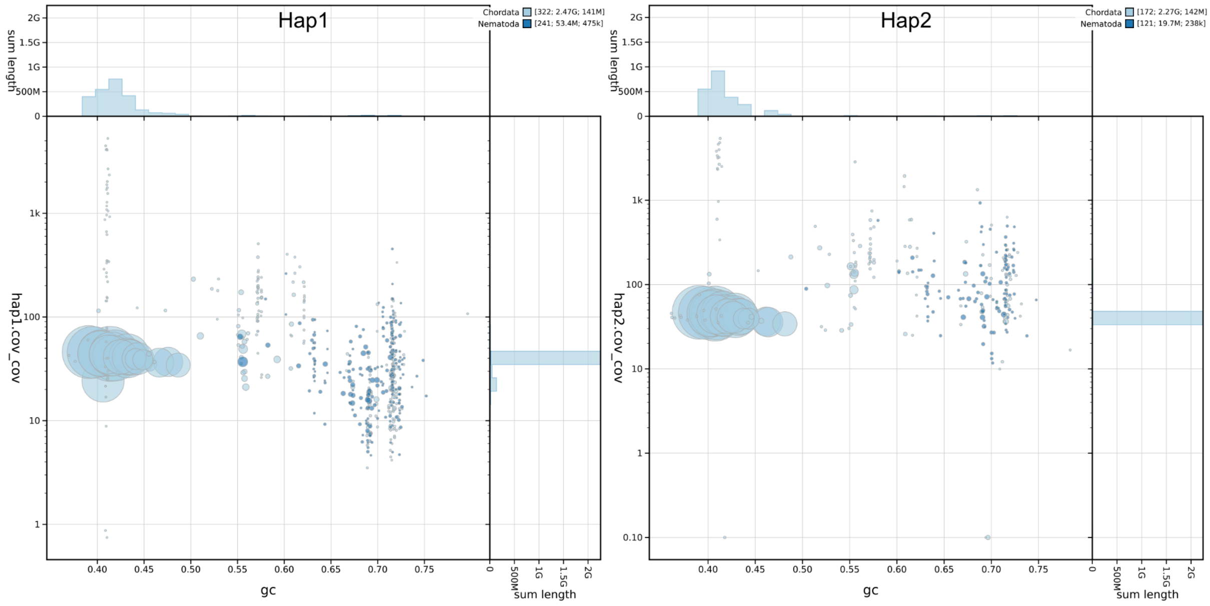

**Supplementary Figure 4: Coverage vs GC plots of *Gulo gulo gulo* hap1 and hap2.** The BlobToolKit blobplots depicts each scaffold as a dot based on the GC content (%GC, x-axis) and coverage (Y-axis). Size of the dots correspond to scaffold length. Dots are colored based on assigned taxonomy. Histograms of sequence lengths within a certain %GC range or coverage range are depicted on the top and right respectively.
